# Structure of the human lipid-sensitive cation channel TRPC3

**DOI:** 10.1101/286153

**Authors:** Chen Fan, Wooyoung Choi, Juan Du, Wei Lü

## Abstract

The TRPC channels are crucially involved in store-operated calcium entry and calcium homeostasis, and they are thus implicated in human diseases such as neurodegenerative disease, cardiac hypertrophy, and spinocerebellar ataxia. We present structure of the full-length human TRPC3, a lipid-gated TRPC member, in a lipid-occupied, closed state at 3.3 Angstrom. TRPC3 has an acorn-like shape with four elbow-like membrane reentrant helices prior to the first transmembrane helix. The TRP helix is perpendicular to, and thus disengaged from, the pore-lining S6, suggesting a different gating mechanism. The third transmembrane helix S3 is remarkably long, resulting in a windmill-like transmembrane domain, and constituting an extracellular domain that may serve as a sensor of external stimuli. We identified two lipid binding sites, one being sandwiched between the pre-S1 elbow and the S4-S5 linker, and the other being close to the ion-conducting pore, where the conserved LWF motif of the TRPC family is located.

## Introduction

The cytosolic free Ca^2+^ concentration is strictly regulated because calcium is crucial to most cellular processes, from transcription control, to neurotransmitter release, to hormone molecule synthesis (Berridge et al., 2003; Kumar and Thompson, 2011; Sudhof, 2012). A major mechanism regulating calcium homeostasis is store-operated calcium entry (SOCE), which is triggered by the depletion of calcium stored in the endoplasmic reticulum (ER) (Ong et al., 2016; Smyth et al., 2010). This process activates store-operated channels (SOCs) in the plasma membrane, resulting in the influx of calcium that refills the calcium stores of the ER for further cellular stimulation (Prakriya and Lewis, 2015). A key component of SOCE has been identified as the TRPC channels, which are calcium-permeable, nonselective cation channels belonging to the TRP superfamily (Liu et al., 2003; Zhu et al., 1998; Zhu et al., 1996).

Among the seven members in TRPC family, TRPC3, TRPC6, and TRPC7 are the closest homologues, and they are unique in being activated by the lipid secondary messenger diacylglycerol (DAG), a degradation product of the signaling lipid phosphatidylinositol 4,5-bisphosphate (PIP2) (Itsuki et al., 2012; Tang et al., 2001). However, the molecular mechanism of such activation remains elusive due to a lack of knowledge of the lipid binding sites. TPRC3, TRPC6, and TRPC7 share several functional domains, including N-terminal ankyrin repeats (AR), a transmembrane domain (TMD) with six transmembrane helixes (S1-S6), and a C-terminal coiled-coil domain (CTD). They also exhibit an unusually long S3 helix, but the function of the S3 helix is poorly understood (Vazquez et al., 2004).

TRPC3 is abundantly expressed in the cerebellum, cerebrum, and smooth muscles, and it plays essential role in the regulation of neurogenesis and extracellular/intracellular calcium signaling (Gonzalez-Cobos and Trebak, 2010; Li et al., 1999). Dysfunction of TRPC3 has been linked to neurodegenerative disease, cardiac hypertrophy, and ovarian adenocarcinoma (Becker et al., 2011; Kitajima et al., 2016; Yang et al., 2009). Although TRPC3 has wide pharmaceutical applications in treatment of these diseases, drug development specifically targeting TRPC3 has been limited due to the lack of understanding of its molecular activation mechanisms (Oda et al., 2017; Xia et al., 2015). Here we report the structure of full-length human TRPC3 (hTRPC3) in a lipid-occupied, inactive state at an atomic resolution of 3.3 Å using single-particle cryo-electron microscopy (cryo-EM). Our structure revealed the first atomic view of TRPC3 channel and its two lipid binding sites, providing insight into the mechanisms of lipid activation and regulation of Ca^2+^ homeostasis.

## Overall architecture

Our three-dimensional reconstruction of hTRPC3 was of sufficient quality to allow *de novo* modeling of almost the entire protein (Fig. S 1, 2), with the exception of the first 21 N-terminal residues; the region connecting the TRP helix and the C-terminal domain (residues 688-757); the loop connecting the linker domain LD6 and LD7 (residues 281-291); and the last 30 C-terminal residues. Characterized by a distinctive one head–two tails shape, we identified two lipid-like densities, one sandwiched between the pre-S1 elbow and the S4-S5 linker, and the other wedged between the P loop and S6 of the adjacent subunit. Notably, we modeled two lipid molecules at these two sites, nevertheless, we were not able to determine the identity of the lipids at current resolution. Interestingly, the TRP helix is perpendicular to the S6, and the density of the hinge region is poorly defined, even though both the TRP helix and S6 exhibit excellent densities (Figure 1a, b).

**Figure 1.**
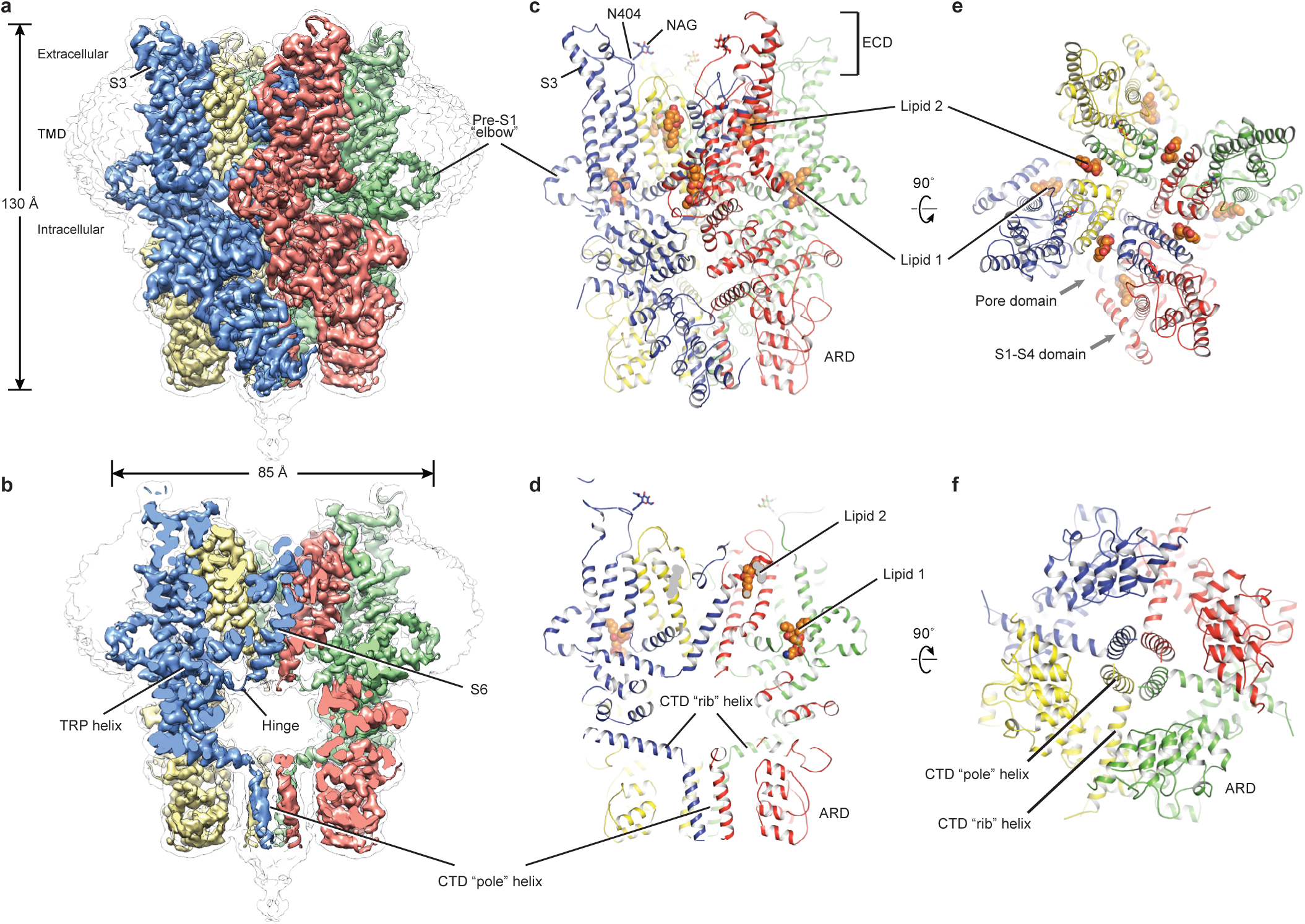
Architecture of human TRPC3. (a) Three-dimensional reconstruction viewed parallel to the membrane. The transparent envelope denotes the unsharpened reconstruction. (b) Slice view of the reconstruction showing the interior of the channel. (c-f) Atomic model of TRPC3 viewed parallel to the membrane (c-d), from the extracellular side (e), and from the intracellular side (f). Each subunit is colored differently.

The overall structure of TRPC3 resembles an acorn, and it has a solely alpha-helical composition (Figure 1a-d). While TRPC3 shares a similar architecture of the TMD with other TRPCs, the third transmembrane helix, S3, is nearly twice as long as the S3 in any other DAG-insensitive TRPC channels (Fig. S 3). It elongates into the extracellular space and connects to the S4 through a remarkably long loop, where a glycosylation site is observed (Figure 1c, d). The extended structure of S3 gives rise to a windmill-like TMD, distinctive to voltage-gated potassium channels or TRP channels (Guo et al., 2017; Long et al., 2007; Paulsen et al., 2016; Shen et al., 2016; Winkler et al., 2017) (Figure 1e). Four elbow-like pre-S1 domains extrude from the TMD and are completely buried in detergent micelles, where the lipid 1 density is located (Figure 1, 2b). The C-terminal coiled-coil domain (CTD) is reminiscent of the TRPM4 and TRPM8 structures, having a coiled-coil “pole” domain in the four-fold symmetry axis, and the “rib” helix penetrating into a “tunnel” composed by adjacent intracellular domains (Figure 1f). This structure thus stabilizes the tetrameric assembly through hydrophobic and polar interactions (Figure 1b, 2b) (Winkler et al., 2017; Yin et al., 2018). The ankyrin repeat domain (ARD), located on the bottom of the channel and comprising four pairs of ARs, is significantly smaller than the ARD of TRPA1 and NOMPC (Jin et al., 2017; Paulsen et al., 2016) (Figure 2b, c).

**Figure 2.**
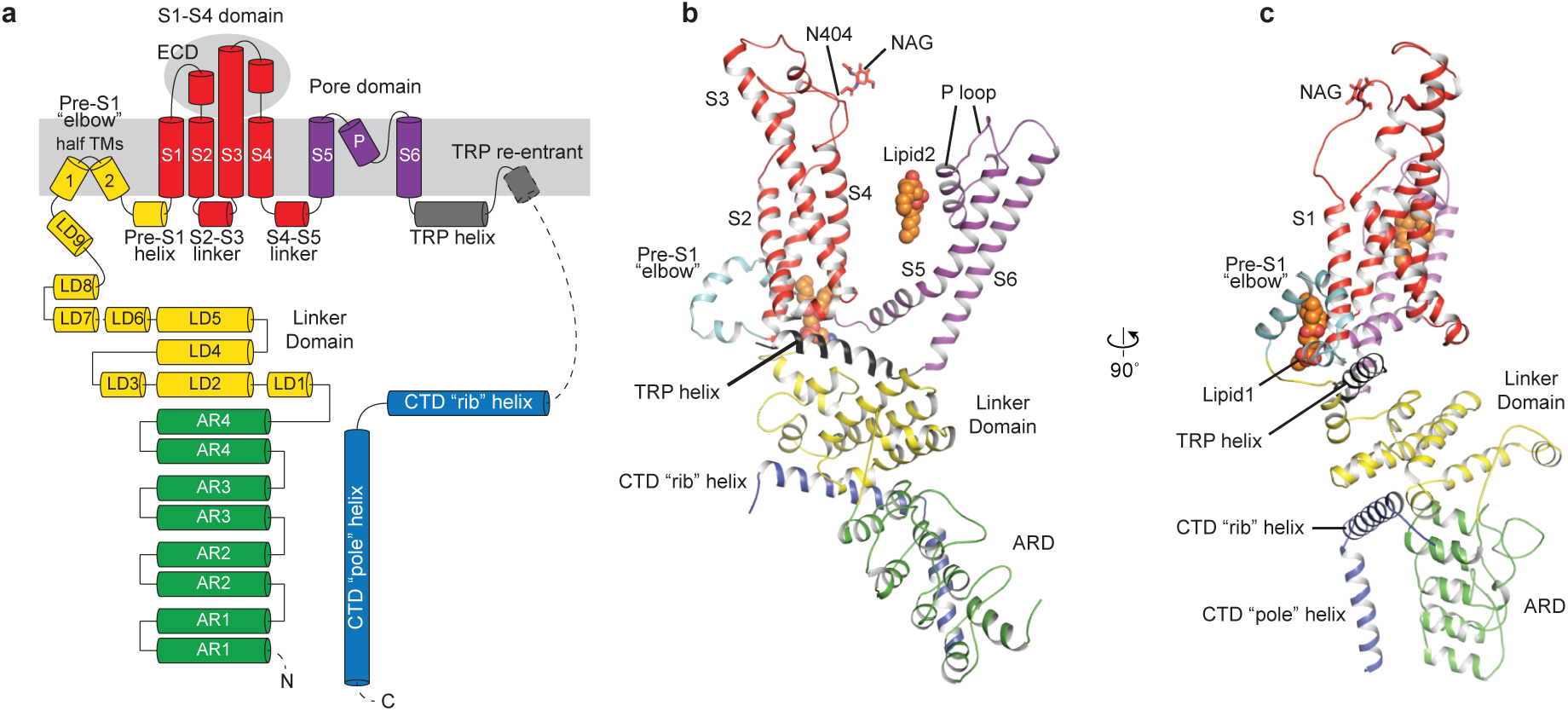
Structure of a single subunit. (a) The schematic representation of TRPC3 domain organization. Dashed lines indicate the regions that have not been modeled. (b-c) Cartoon representation of one subunit color-coded to match panel a.

## Transmembrane domain and lipid-binding sites

The TMD of TRPC3 shares topology similar to that of other TRP channels and voltage-gated ion channels, consisting of the S1-S4 domain and the pore domain arranged in a domain-swapped manner (Figure 1e, 3a). Nevertheless, the distinct activation mechanism of TRC3, TRPC6, and TRPC7 by DAG implies unique features of their TMD. Indeed, comparison of the relative arrangement of the S1-S4 domains with the pore domain shows remarkable differences between TRPC3 and TRPA1 or TRPM4, yet overall agreement with TRPV1 (Fig. S 4). Detailed inspection of the TMD in TRPC3 reveals two unique features: a large elbow-like pre-S1 domain harboring a lipid-binding site (lipid 1), and unusually long S3 helix forming an extracellular domain (ECD), along with the S1-S2 linker and S3-S4 linker (Figure 3a, b).

**Figure 3.**
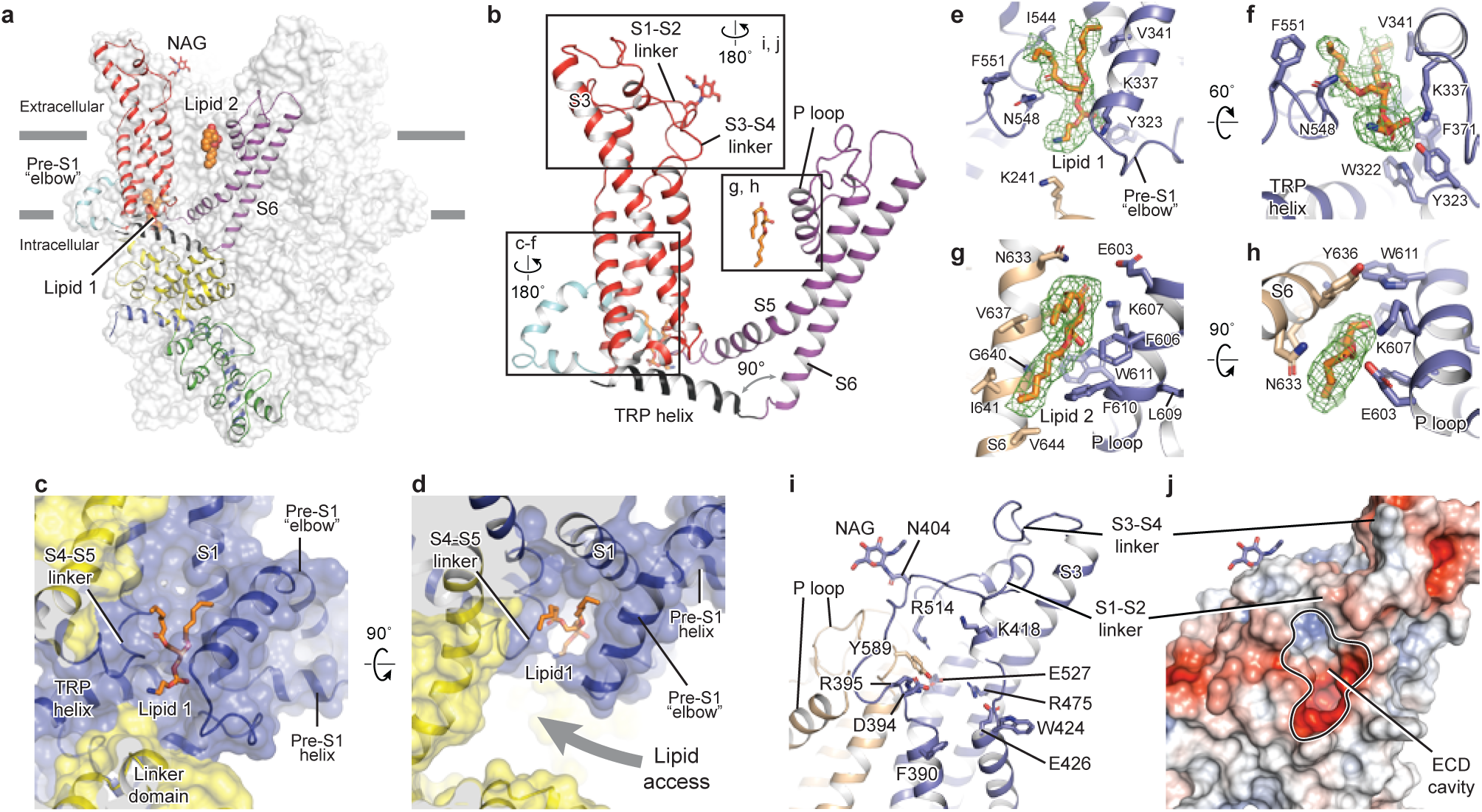
Transmembrane domain, extracellular domain, and lipid-binding sites. (a) Domain organization. The channel is shown in surface representation, with one subunit shown in cartoon representation. The colors match those in Figure 2a. (b) Details of the transmembrane domain and extracellular domain. (c-d) Pre-S1 elbow and binding site of lipid 1. The lipid molecule is buried inside the pocket formed by pre-S1 elbow, S1, and the S4-S5 linker. Two adjacent subunits (blue and yellow) are shown in both cartoon and surface representations. The lipid molecule is shown as sticks. (e-f) Residues that interact with lipid 1 are shown in sticks, and protein is shown in cartoon representation. Lipid density is shown in mesh. (g-h) Lipid 2 binds between S6 and the P loop of adjacent subunits, which are in light blue and wheat. (i) Structure of the ECD. Key residues forming the cavity is shown in sticks. Adjacent subunits are in light blue and wheat. (j) Surface representation of the ECD, colored according to the electrostatic surface potential. The color gradient is from -5 to 5 kT/*e* (red to blue).

The pre-S1 elbow, embedded in the lipid bilayer, consists of two half transmembrane helices (half TM1 and half TM2). The half TM1 connects to LD9, which is the last alpha helix in the LD; the half TM2 connects to the pre-S1 helix, a short alpha helix prior to S1 running horizontally along the intracellular face of the membrane (Figure 2a, 3c, 3d). This unique configuration pulls the intracellular half of S1 away from the pore center, resulting in a hydrophobic pocket behind the pre-S1 elbow and surrounded by half TM1 and S1 (Figure 3c). Moreover, the outward movement of S1 opens a window between itself and S5 from the adjacent subunit, exposing the intracellular half of S4 and the S4-S5 linker, which are key regions for TRP channel gating, to the lipid environment (Figure 3d).

Indeed, we observed a lipid-shaped density (lipid 1) in this pocket (Figure 3c, d). The head group of lipid 1 is well defined in the density map, forming several hydrogen bonds and polar interactions with residues in the LD9, the pre-S1 elbow, half TM1, and the S4-S5 linker, while the two hydrocarbon tails are in contact with S1, S4, the pre-S1 elbow, and half TM1 (Figure 3e, f). A similar pre-S1 elbow structure with lipid-like density has been observed in the *Drosophila* mechanosensitive channel NOMPC (Jin et al., 2017). We suggest that this lipid site may be crucially linked to channel activation, given its interaction with S4 and the S4-S5 linker. A mutation in this region (T561A on S4) results in gain of function, causing abnormal Purkinje cell development and cerebellar ataxia in moonwalker mice (Becker, 2014) (Fig. S 3).

We also identified a second lipid-like density (lipid 2) in the lateral fenestration of the pore domain, wedged between the P loop and S6 of adjacent subunit and forming both hydrophobic and hydrophilic interactions (Figure 3g, h). Moreover, lipid 2 is in close contact with the LFW motif on the P loop, which is highly conserved throughout the TRPC family and is crucial to channel function (Fig. S 3). Replacing this motif by three alanine residues in TRPC5 and TRPC6 resulted in a nonfunctional channel (Strubing et al., 2003). Therefore, the lipid 2 binding site likely represents another important modulation site. In addition to interaction with lipid 2, the LFW motif forms multiple hydrophobic interactions within the pore domain and therefore plays an important role in maintaining the proper structure of the pore domain (Figure 3i).

A second unique feature of TRPC3 is the remarkably long S3, stretching out into the extracellular side and supporting the formation of the ECD (Figure 3a). Within the ECD we observed a cavity-like feature (Figure 3j), with S3 and the S3-S4 linker as a “back wall and roof”, and the S1-S2 linker forming the entrance. This cavity is located right above the lipid bilayer, and its interior is filled with both charged and hydrophobic residues (Figure 3j). Moreover, a tyrosine residue (Y589) in the loop connecting the S5 and the P loop plugs into the cavity (Figure 3i). We speculate that the cavity may serve as a binding site for small molecules and that binding of small molecules may directly affect channel function through Y589, implying a role for the ECD as a sensor of external stimuli. This is in line with the finding that Pyr3, a TRPC3-specific inhibitor, likely binds to the extracellular side of the protein (Kiyonaka et al., 2009). Furthermore, a glycosylation site (N404) is observed in the S1-S2 loop, consistent with the prediction that TRPC3 is monoglycosylated in the extracellular side (Figure 3i) (Vannier et al., 1998). The site is very close to the P loop, suggesting that the glycosylation status may affect channel activity, and this is consistent with the report that N-linked glycosylation is a key determinant of the basal activity of TRPC3 (Dietrich et al., 2003). Further studies are necessary to clarify the physiological role of the ECD.

## TRP domain

The TRP domain—the namesake region in the TRP channel located at the border between the transmembrane domain and the intracellular domain—is crucially involved in signal transduction and channel gating (Garcia-Sanz et al., 2007; Taberner et al., 2013). Similar to that of TRPM4, the TRP domain consists of a TRP helix that runs nearly parallel along the intracellular face of the membrane and a TRP re-entrant helix embedded in the lipid bilayer (Autzen et al., 2018; Guo et al., 2017; Winkler et al., 2017) (Figure 4a). The TRP helix penetrates into the tunnel formed by the S4-S5 linker of the TMD on the top and the LD9 of the linker domain in the intracellular space on the bottom (Figure 4a), showing an apparently disengaged connection to the S6 helix through a loop of the hinge region instead of a continuous alpha helical structure as in TRPM4 (Figure 4b, e). While the densities for both S6 and TRP helix were well defined, their linker region was surprisingly poorly defined, indicating a high flexibility between the TRP helix and S6 (Figure 4b, 1b). The TRP helix forms an approximate right angle to the S6, in strong contrast to the TRPV1, TRPA1, and TRPM4 structures whose TRP helices form obtuse angles with S6 (Figure 4a-d). Given the crucial role of TRP helix in channel gating and its possible involvement in voltage dependence (Nilius et al., 2005b), the disengagement of TRP helix from the pore-lining S6 may provide a molecular basis for the unique gating mechanism of TRPC3 relative to other TRP subfamily channels (Itsuki et al., 2012).

**Figure 4.**
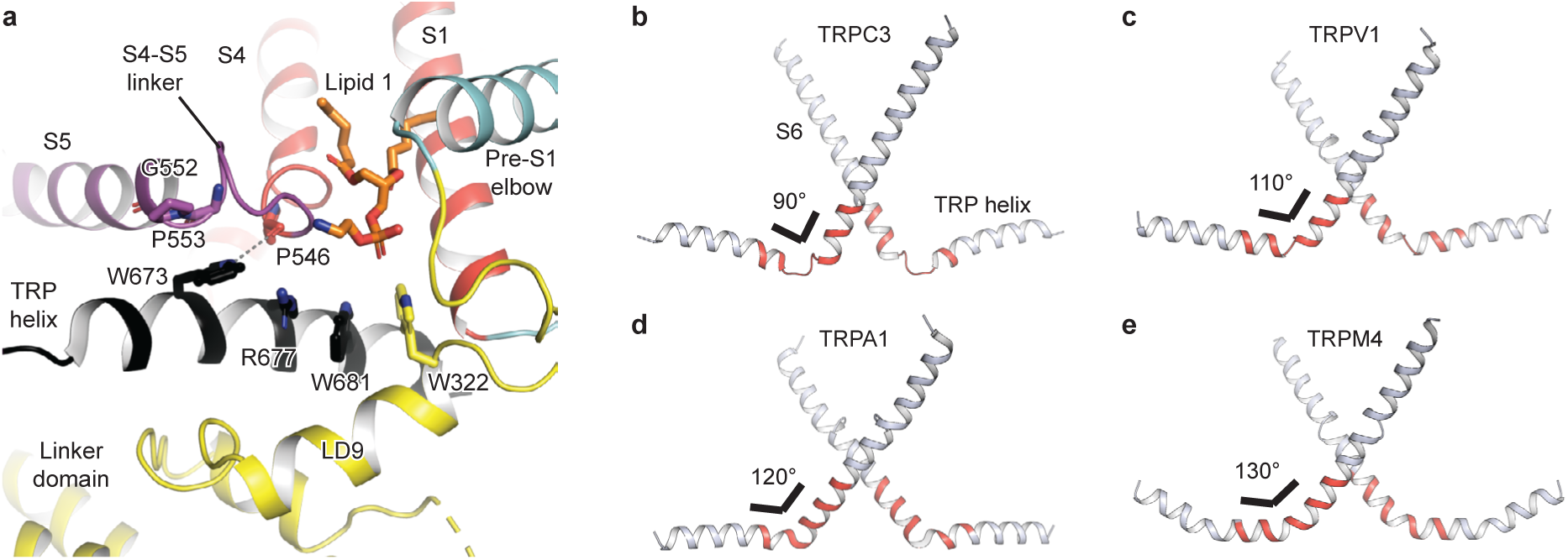
The TRP domain. (a). Cartoon representation of the TRP helix, pre-S1 elbow, TMD, and linker domain, showing their interaction. Lipid 1 is shown in sticks. W673 on the TRP helix stacks with P553 and G552 forming a hydrogen bond with the backbone oxygen (dashed line) of P546 on the S4-S5 linker. The side chain of R677 is in close contact with the head group of lipid 1. (b-e). The pore lining helix S6 and the TRP helix in TRPC3 (b), TRPV1 (c), TRPA1 (d), and TRPM4 (e). The angle between the S6 and TRP helices are indicated; only two subunits are shown for clarity. The hinge connecting the S6 and TRP helix is highlighted in red.

Furthermore, the TRP helix forms a series of polar and hydrophobic interactions with the S4-S5 linker and the LD9 helix (Figure 4a). Specifically, the highly conserved tryptophan W673 is extensively coupled with the S4-S5 linker through interactions with G552, P553, and P546. Mutation of the corresponding W673 in TRPV3 results in Olmsted syndrome (Ni et al., 2016), and replacement of the corresponding residue in NOMPC results in a channel that has increased current amplitude but is nonresponsive to mechanical stimuli(Jin et al., 2017). Mutation of the corresponding tryptophan in TRPV1 abolishes channel activation in response to depolarization (Gregorio-Teruel et al., 2014). Replacement of the corresponding G552 in TRPC4 and TRPC5 by serine results in a constantly open channel (Beck et al., 2013). Another highly conserved tryptophan residue in the TRP helix, W681, tightly packs with W322 in the LD9 (Figure 4a and Fig. S 3). Interestingly, the highly conserved R677 in the TRP helix is close to the head group of lipid 1, and its replacement by histidine increases channel activity and results in neuronal cell death and cerebellar ataxia, perhaps by affecting the binding of lipid 1 (Figure 4a) (Fogel et al., 2016).

## Ion-conducting pore

The ion-conducting pore of TRPC3 is lined with an extracellular selectivity filter and an intracellular gate, with a wide central vestibule in the middle (Figure 5a). The pore adopts a closed conformation with the narrowest radius - at I658 and L654 on S6 close to the intracellular exit - of less than 1 Å, thus preventing ion passage (Figure 5b). Presumably, the channel is trapped in a lipid-bound inactive state or the bound lipids are not the activator DAG. The selectivity filter is defined by the backbone carbonyl oxygens of I613, F614 and G615 located in the P loop. The narrowest point at G615 has a radius of 2.1 Å, allowing partially dehydrated ions to pass through (Figure 5b, c). Moreover, five acidic residues in the P loop and the intracellular end of S6 in TRPC3 impose a negative electrostatic surface potential, which is important for cation selectivity (Figure 5a, d). On the intracellular site, the inner surface along the CTD and ARD contains acidic residues, giving rise to a negative charge and thus providing a possible pathway by which cations can access the cytoplasm (Figure 5a, e).

**Figure 5:**
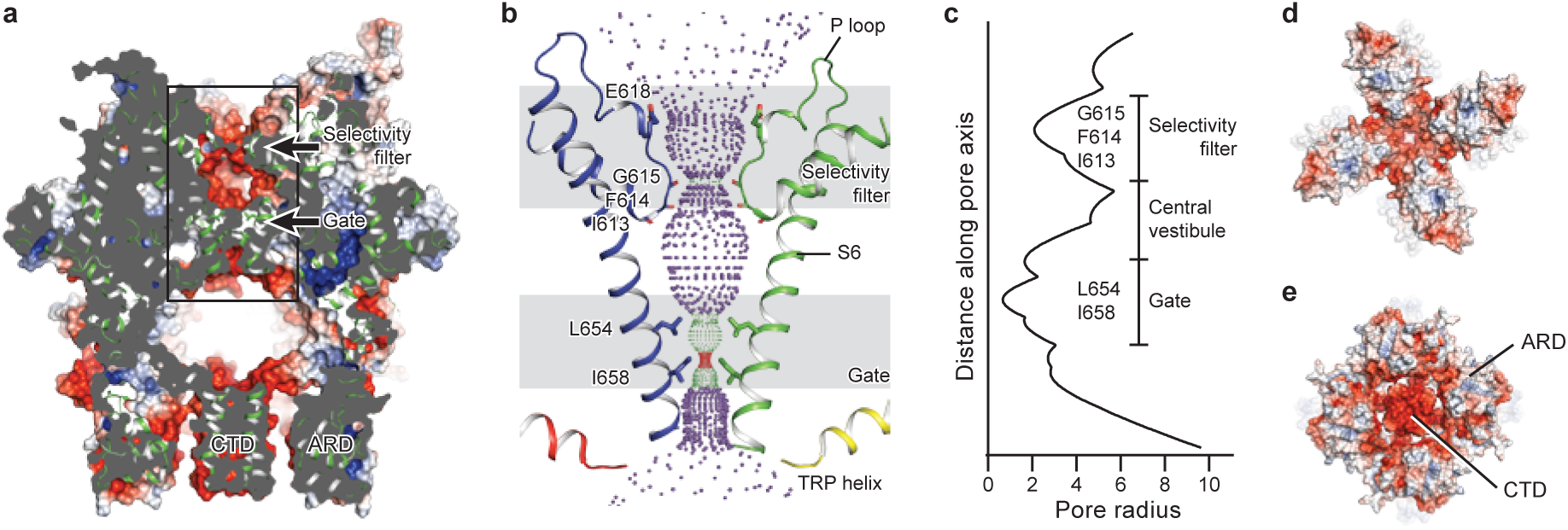
The ion-conducting pore. (a, d, e) Surface representation of TRPC3, viewed (a) parallel to the membrane, (d) from the extracellular side, and (e) from the intracellular side. The surface is colored according to electrostatic surface potential; the color gradient is from -5 to 5 kT/*e* (red to blue). The protein is also shown in cartoon representation in (a). (b) The shape and size of the ion-conducting pore (boxed area in panel a). The P loop and S6 of two subunits and the TRP helix of the other two subunits are shown as cartoons, and the side chains of restriction residues are shown as sticks. Purple, green, and red spheres define radii of > 2.3, 1.2–2.3, and < 1.2 Å, respectively. (c) Plot of pore radius as a function of distance along the pore axis in Angstroms.

Similar to other Ca^2+^-permeable TRP channels, an acidic residue, E618, is located at the entrance of the selectivity filter. An E618Q mutation impedes the calcium permeability of TRPC3, but it preserves monovalent permeation (Feng et al., 2013; Liu et al., 2007) (Figure 5b). The neutralization of the corresponding acidic amino acid on TRPV1 remarkably decreases channel’s permeability to divalent ions (Garcia-Martinez et al., 2000). By contrast, replacement of a glutamine residue at the corresponding position (Q977) by an acidic amino acid in TRPM4, which is a Ca^2+^-impermeable TRP channel, produced moderate Ca^2+^ permeability (Nilius et al., 2005a). Thus, having an acidic residue close to the selectivity filter may represent a general principle of permeability for divalent cations in nonselective Ca^2+^-permeable TRP channels.

## The intracellular domain

TRPC3 exhibits a similar intracellular domain composition as TRPA1, including a C-terminal CTD and a N-terminal ARD. We found several unanticipated features that advance our understanding of the molecular basis of TRPC family (Figure 6a). First, the ARD in TRPC3, consisting of 4 ARs, is significantly shorter than that in TRPA1 (16 repeats). Second, instead of a straight coiled-coil domain as in TRPA1, TRPC3 adopts the characteristic umbrella-like CTD “pole” and “rib” domain of the TRPM family (Figure 6b, c). Interestingly, the turn from the pole to the rib helix is where the ankyrin repeats end. Third, between the rib domain and TRP helix, there is a linker domain that has remarkable structural similarities to the MHR4 (TRPM homology region) domain in TRPM4 (Autzen et al., 2018; Guo et al., 2017; Winkler et al., 2017), as well as to the linker domain in NOMPC (Jin et al., 2017). The location of the linker domain suggests a role for signal transduction from ARD and CTD further to TMD. Overall, the TRPC3 forms a unique intracellular domain that has structural features characteristic of the TRPM, TRPA, and NOMPC families. Although the functional role of the intracellular domain is yet unknown, it clearly contributes to the channel assembly through three major interfaces. The first interface is contributed by the vertical CTD pole helices of the four subunits winding into a tetrameric coiled-coil assembly (Figure 6b, c). This is a common feature employed to specify subunit assembly and assembly specificity within the voltage-gated ion channel superfamily (Figure 6b, c). The second interface is formed by the horizontal CTD rib helix penetrating through the tunnel composed of ARD and LD from neighboring subunits, thus tethering them together (Figure 6c, d). Notably, the rib helix is rich in positively charged residues, forming multiple interactions with the charged residues in the LD. The third interface is located between LD and LD/pre-S1 elbow of the adjacent subunit (Figure 6e). All these interactions knit the tetramer together.

**Figure 6.**
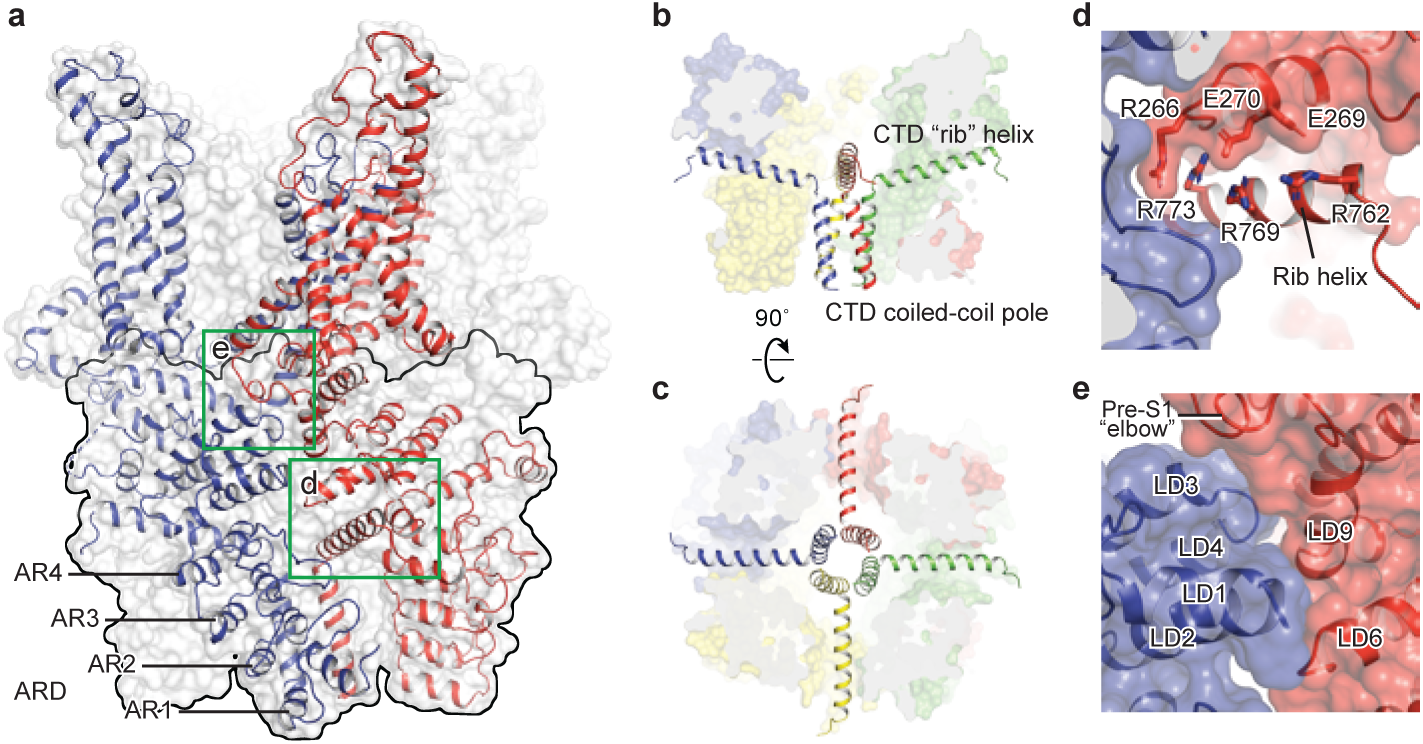
The intracellular domain. (a) Surface representation of TRPC3 with two adjacent subunits shown in cartoon representation. The intracellular domain is highlighted in the black frame. Two interfaces highlighted in green frames are enlarged in (d-e). (b-c) Cartoon representation of the CTD coiled-coil pole and the rib helix. The intracellular domain is shown in surface representation, viewed in parallel to the membrane (b) and from the intracellular side (c). (d) Inter-subunit interface formed by the CTD rib helix with adjacent ARD and LD. Protein is shown in cartoon and surface representations. Two adjacent subunits are in blue and red. Charged residues forming hydrogen bond or polar interaction with each other are shown as sticks. (e) Interface between adjacent LDs and pre-S1. Alpha helices involved in the inter-subunit interaction are indicated.

## Conclusion

The TRPC3 structure displays a unique acorn-like architecture. Distinct to the TRPM, TRPV or TRPA channels whose TRP helix and S6 form a continuous alpha helical structure, the TRP helix in TRPC3 is disengaged from the S6, which aligns with the unique gating mechanism of TRPC, perhaps linked to the lipid activation or voltage independence. The remarkably long S3 endows TRPC3 a windmill-like TMD and frames the ECD in which a cavity may act as a binding site for small molecules, suggesting a role for the ECD in sensing extracellular stimuli. We identified two lipid binding sites, one buried in a pocket surrounded by the pre-S1 elbow, S1, and the S4-S5 linker, and the other inserted into the lateral fenestration of the pore domain. Our structure provides a framework for understanding the complex gating mechanism of TRPC3.

---

**Fig. S 1.**
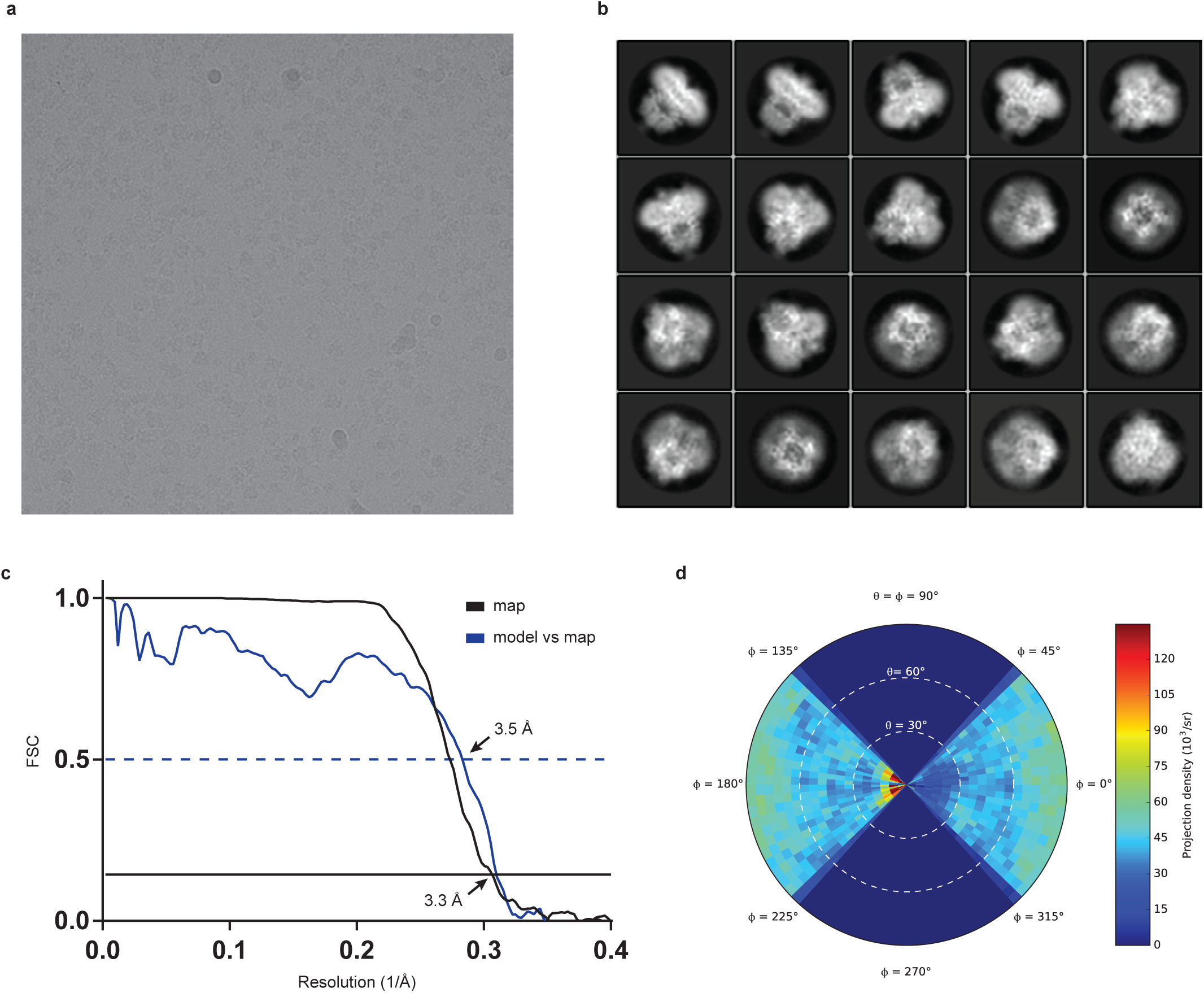
Cryo-EM analysis of human full-length TRPC3. (a) Representative electron micrograph. (b) Selected two-dimensional class averages of the electron micrographs. (c) The gold-standard Fourier shell correlation curve for the EM maps is shown in black and the FSC curve between the atomic model and the final EM map is shown in blue. (d) Angular distribution of particles used for refinement.

**Fig. S 2.**
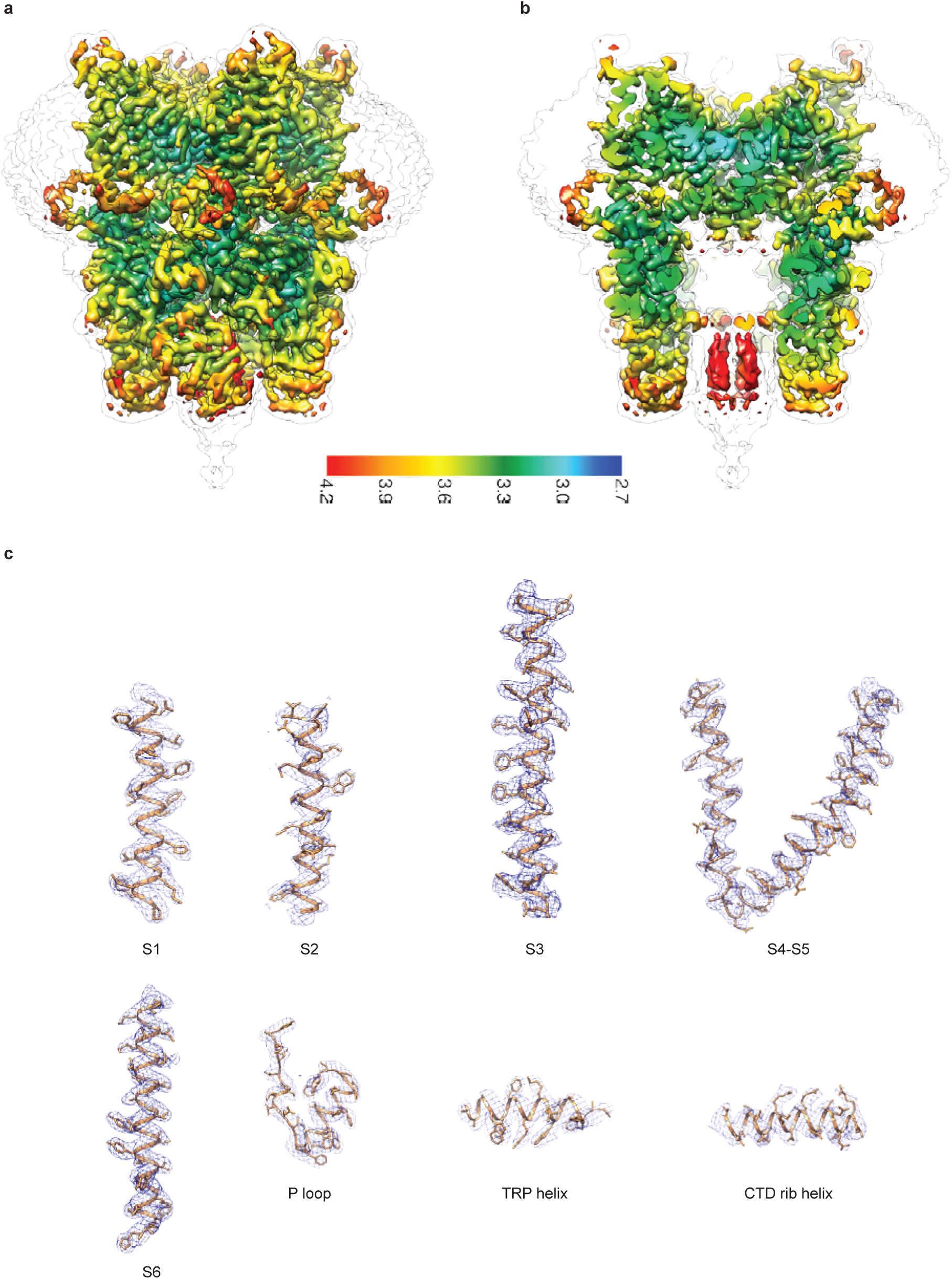
Cryo-EM map of human full-length TRPC3. (a-b) Local resolution estimation. The map is colored according to local resolution estimation. (c) Representative densities Density maps are shown in blue meshes, and the atomic models are shown in cartoon representation with side chains as sticks.

**Fig. S 3.**
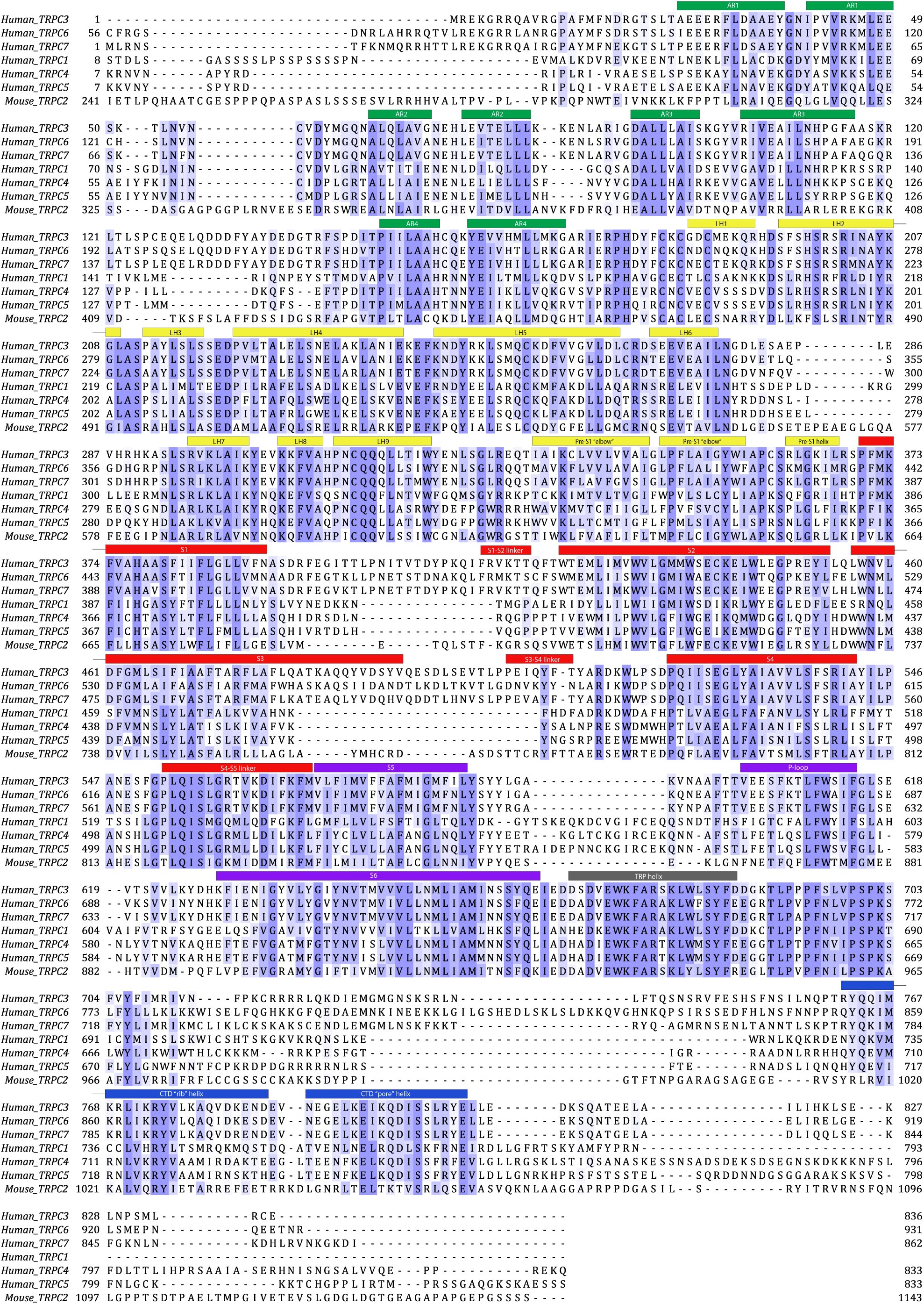
Secondary structure arrangement of human TRPC3 and sequence alignment of TRPC family channels. The TRPC2 is from *mus musculus*, whereas all the other proteins are from human. The sequences were aligned using the Clustal Omega program on the Uniprot website and coloured using BLOSUM62 score by conservation. The secondary structural elements are color-coded to match Figure 2a.

**Fig. S 4.**
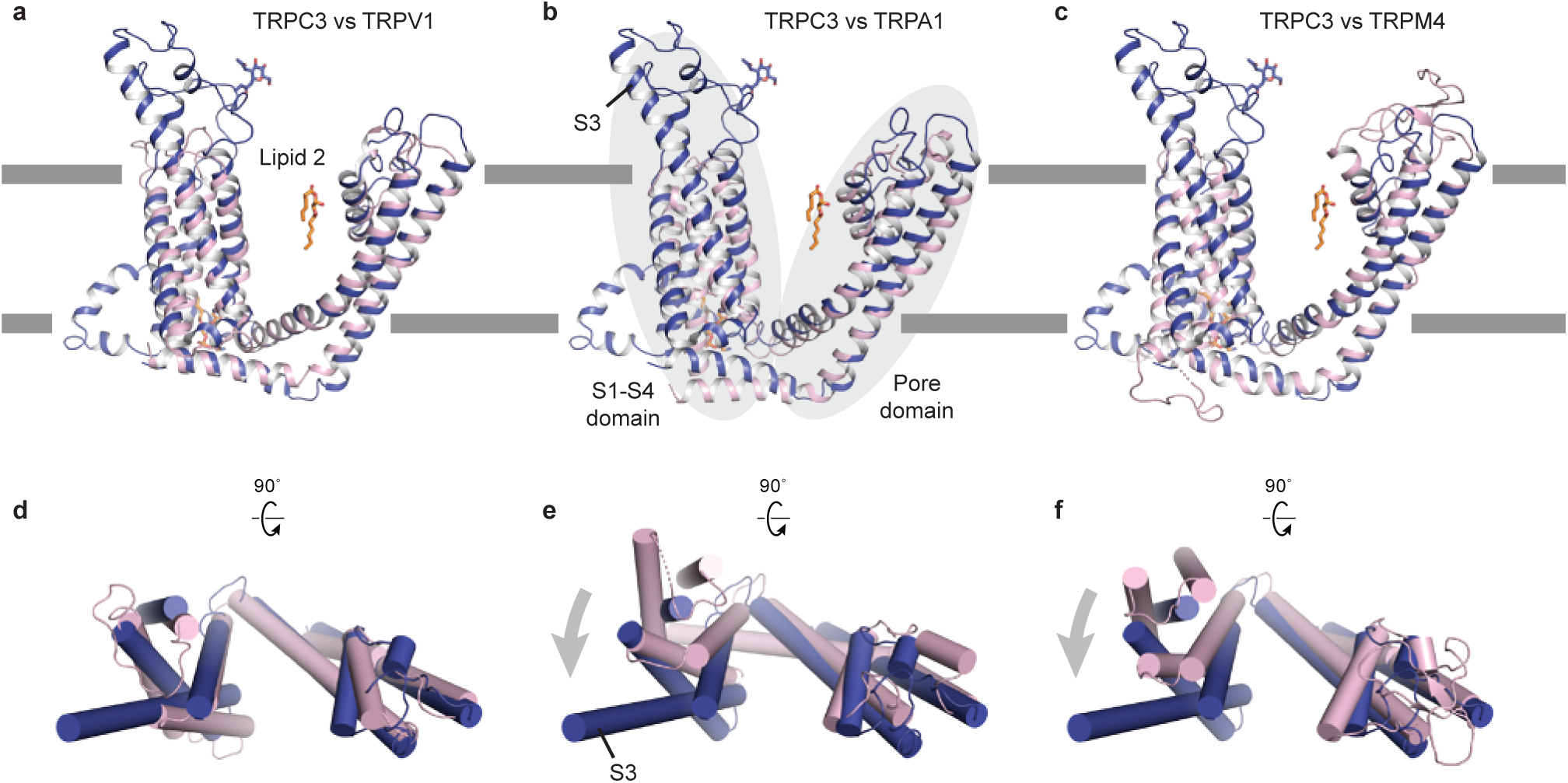
Comparison of the TMD of TRPC3 with TRPV1 (a), TRPA1 (b), and TRPM4 (c). Structures are aligned using main chain atoms of the pore domain. Only the TMD of one subunit is shown in cartoon representation, viewed in parallel to the membrane. TRPC3 is in blue; TRPV1, TRPA1, and TRPM4 are in pink. (d-e) TMD viewed from extracellular side. The relative organization of the S1-S4 domain with the pore domain in TRPC3 is similar to that in TRPV1, but the S1-S4 domain in TRPC3 exhibits a clockwise rotation relative to TRPA1 or TRPM4.

**Table S 1.** Statistics of EM data processing and model refinement.

|  |  |
| --- | --- |
| **Data collection/processing** |  |
| Microscope | Titan Krios (FEI) |
| Voltage (kV) | 300 |
| Defocus range ($\mu\text{M}$) | 1.0 – 2.5 |
| Exposure time (s) | 8 |
| Dose rate ($e^-/\text{\AA}^2/\text{s}$) | 6.76 |
| Number of frames | 40 |
| Pixel size ($\text{\AA}$) | 1.074 |
| Particles refined | 143855 |
| Resolution ($\text{\AA}$) | 3.3 |
| FSC threshold | 0.143 |
| Resolution range ($\text{\AA}$) | 412.4 – 3.3 |
| **Model statistics** |  |
| Number of atoms | 20988 |
| Protein | 20744 |
| Ligand | 244 |
| r.m.s. deviations |  |
| Bond length ($\text{\AA}$) | 0.005 |
| Bond angle ($^\circ$) | 1.008 |
| Ramachandran plot |  |
| Favored (%) | 94.09 |
| Allowed (%) | 5.77 |
| Disallowed (%) | 0.14 |
| Rotamer outlier (%) | 0.85 |
| Clashscore | 3.0 |

## Methods

### Construct, expression and purification of TRPC3

A full-length human *TRPC3* gene (UniProtKB (http://www.uniprot.org) accession number, Q13507) with 836 amino acid residues was synthesized by Genscript and was subcloned into a modified version of pEG BacMam vector containing: a twin strep-tag, a His8-tag, and green fluorescent protein (GFP) with thrombin cleavage site at the N terminus (Goehring et al., 2014). The recombinant Bacmid DNA and baculovirus of TPRC3 were generated by sf9 insect cells, and P2 viruses were used to infect suspension HEK293 cells.

For large-scale expression, suspension HEK293 cells were cultured in Freestyle 293 expression Medium (Invitrogen) with 1% (v/v) fetal bovine serum (FBS). When cell density reached around 3 million/ml, 8% (v/v) of P2 viruses were introduced. At 12 h post-infection, 10 mM sodium butyrate was supplemented and then cells were transferred to 30 °C. The infected cells were collected at 48 h post-infection by centrifugation at 4000 rpm for 15 min at 4 °C and then was washed once with TBS buffer (20 mM Tris, pH 8.0, 150 mM NaCl).

TRPC3 was extracted from the cells by solubilization buffer containing 20 mM Tris 8.0, 500 mM NaCl in the presence of 1 mM PMSF, 0.8 μM aprotinin, 2 μg/ml leupeptin, and 2 mM pepstatin A with 1% digitonin (Calbiochem) for 2 h at 4 °C. The cell debris were eradicated by ultracentrifugation at 40,000 rpm using 45 Ti rotor (Beckman Coulter, Inc.) for 1 h at 4 °C. The solubilized proteins were incubated with TALON resin and the resin was washed with 10 column volumes of wash buffer (20 mMTris 8.0, 500 mM NaCl, 15 mM imidazole, and 0.1% digitonin). The TALON resin-bound TRPC3 was eluted with elution buffer (20 mM Tris 8.0, 500 mM NaCl, 250 mM imidazole, and 0.1% digitonin). Thrombin (1:20 molar ratio) and 10 mM EDTA were added into the eluted sample and incubated for 3 h on the ice. In order to further purify the protein, the sample was concentrated and loaded onto a superpose6 column in buffer containing 20 mM Tris 8.0, 500 mM NaCl, 1 mM EDTA with 0.1% digitonin. Peak fractions containing TRPC3 were pooled and concentrated to 5 mg/ml.

### EM sample preparation and data acquisition

The purified TRPC3 protein sample (2.5 μL) at a concentration of 5 mg/mL was applied onto a glow-discharged Quantifoil holey carbon grid (gold, 1.2/1.3 μm size/hole space, 300 mesh). The gird was blotted for 1.5 s at 100% humidity by using a Vitrobot Mark III, and then was plunged into liquid ethane cooled by liquid nitrogen. Images were obtained by an FEI Titan Krios electron microscope operating at 300 kV with a nominal magnification of 130,000× Gatan K2 Summit direct electron detector was used in order to record image stacks in super-resolution counting mode with a binned pixel size of 1.074 Å. Every image was dose-fractionated to 40 frames with a total exposure time of 8 s with 0.2 s per frame. Dose rate was 6.76 e^−^ Å^−2^ s^−1^. The images stacks were recorded using the automated acquisition program SerialEM (Mastronarde, 2005). Nominal defocus values varied from 1.0 to 2.5 μm.

### EM data processing

MortionCor2 was used to implement motion-correction of summed movie stacks(Zheng et al., 2017). Gctf was applied to estimate Defocuse values (Zhang, 2016). Particles were picked from approximately 200 micrographs using Gautomatch (http://www.mrc-lmb.cam.ac.uk/kzhang/Gautomatch/) and subjected to an initial reference-free 2D classification using Relion 2.1(Scheres, 2012). Nine representative 2D class averages were selected as templates for automated particle picking for the entire data set using Gautomatch. The auto-picked particles were visually checked and obvious bad particles were manually removed. The picked particles were cleaned up throughout three rounds of 2D classification. CryoSPARC was applied to obtain an initial model (Punjani et al., 2017). The selected particles after 2D classification were subjected to 3D classification of 5 classes using Relion 2.1, with the initial reconstruction low-pass-filtered to 60 Å as a reference model. Only one out of five classes presented high-resolution features, hence, particles from this class were combined and further refined via Relion 2.1. Particles were further refined using the local refinement from Frealign with C4 symmetry applied and high-resolution limit for particle alignment set to 4.5 Å(Grigorieff, 2016). The resolutions reported are based on the “limiting resolution” procedure in which the resolution during refinement is limited to a lower resolution than the resolution estimated for the final reconstruction. The final resolutions reported in Table supplement 1 are based on the gold standard Fourier shell correlation (FSC) 0.143 criteria. To calculate the FSC plot, a soft mask (4.3 Å extended from the reconstruction with an additional 4.3 Å cosine soft edge, low-pass filtered to 10 Å) was applied to the two half maps.

### Model building

The model of TRPC3 was built in Coot using the TMD domain of TRPM4 structure (PDB 5wp6) as a guide(Emsley et al., 2010). *De novo* building was mainly guided by bulky residues and secondary structure prediction (Fig. S 3). The TRPC3 structure chiefly consists of α helices, which greatly assisted register assignment. In the initial *de novo*-built model, the order and length of the secondary structure features, as well as the positions of bulky residues within each secondary structure feature are in good agreement with the prediction (Fig. S 3). The initial model was then subjected to real space refinement using Phenix.real_space_refine with secondary structure restraints (Afonine et al., 2012). The refined model was manually examined and re-modified via COOT. For validation of refined structure, FSC curves were applied to calculate the difference between the final model and EM map. The geometries of the atomic models were evaluated using MolProbity (Chen et al., 2010). All figures were prepared using UCSF Chimera and Pymol (Schrödinger) (The PyMOL Molecular Graphics System)(Pettersen et al., 2004).

### Data Availability

The cryo-EM density map and coordinate of TRPC3 have been deposited in the Electron Microscopy Data Bank (EMDB) accession number EMD-XXXX, and in the RCSB Protein Data Bank (PDB) accession code XXXX.

## Acknowledgements

We thank G. Zhao and X. Meng for the support with data collection at the David Van Andel Advanced Cryo-Electron Microscopy Suite. We appreciate the VARI High-Performance Computing team for computational support. We thank D. Nadziejka for technical editing, and C-H. Lee for helpful discussion. This work was supported by internal VARI funding.

## Author Contributions

W. L. and J. D. initiated the project. C. F. and W. C. carried out protein purification and initial cryo-EM experiments, including grid preparation and initial grid screening for optimizing grid conditions. All the authors contributed in cryo-EM data collection and processing, structure analysis, and preparation of the manuscript.

## References

Afonine, P.V., Grosse-Kunstleve, R.W., Echols, N., Headd, J.J., Moriarty, N.W., Mustyakimov, M., Terwilliger, T.C., Urzhumtsev, A., Zwart, P.H., and Adams, P.D. (2012). Towards automated crystallographic structure refinement with phenix.refine. Acta Crystallogr D 68, 352–367.

Autzen, H.E., Myasnikov, A.G., Campbell, M.G., Asarnow, D., Julius, D., and Cheng, Y.F. (2018). Structure of the human TRPM4 ion channel in a lipid nanodisc. Science 359, 228–232.

Beck, A., Speicher, T., Stoerger, C., Sell, T., Dettmer, V., Jusoh, S.A., Abdulmughni, A., Cavalie, A., Philipp, S.E., Zhu, M.X., et al. (2013). Conserved Gating Elements in TRPC4 and TRPC5 Channels. J Biol Chem 288, 19471–19483.

Becker, E.B.E. (2014). The Moonwalker Mouse: New Insights into TRPC3 Function, Cerebellar Development, and Ataxia. Cerebellum 13, 628–636.

Becker, E.B.E., Fogel, B.L., Rajakulendran, S., Dulneva, A., Hanna, M.G., Perlman, S.L., Geschwind, D.H., and Davies, K.E. (2011). Candidate Screening of the TRPC3 Gene in Cerebellar Ataxia. Cerebellum 10, 296–299.

Berridge, M.J., Bootman, M.D., and Roderick, H.L. (2003). Calcium signalling: Dynamics, homeostasis and remodelling. Nat Rev Mol Cell Bio 4, 517–529.

Chen, V.B., Arendall, W.B., Headd, J.J., Keedy, D.A., Immormino, R.M., Kapral, G.J., Murray, L.W., Richardson, J.S., and Richardson, D.C. (2010). MolProbity: all-atom structure validation for macromolecular crystallography. Acta Crystallogr D 66, 12–21.

Dietrich, A., Schnitzler, M.M.Y., Emmel, J., Kalwa, H., Hofmann, T., and Gudermann, T. (2003). N-linked protein glycosylation is a major determinant for basal TRPC3 and TRPC6 channel activity. J Biol Chem 278, 47842–47852.

Emsley, P., Lohkamp, B., Scott, W.G., and Cowtan, K. (2010). Features and development of Coot. Acta Crystallogr D 66, 486–501.

Feng, S.J., Li, H.Y., Tai, Y.L., Huang, J.B., Su, Y.J., Abramowitz, J., Zhu, M.X., Birnbaumer, L., and Wang, Y.Z. (2013). Canonical transient receptor potential 3 channels regulate mitochondrial calcium uptake. P Natl Acad Sci USA 110, 11011–11016.

Fogel, B., Hanson, S., and Becker, E. (2016). Mutation of the Murine Ataxia Gene TRPC3 Causes Cerebellar Ataxia in Humans. Neurology 86, 284–286.

Garcia-Martinez, C., Morenilla-Palao, C., Planells-Cases, R., Merino, J.M., and Ferrer-Montiel, A. (2000). Identification of an aspartic residue in the P-loop of the vanilloid receptor that modulates pore properties. J Biol Chem 275, 32552–32558.

Garcia-Sanz, N., Valente, P., Gomis, A., Fernandez-Carvajal, A., Fernandez-Ballester, G., Viana, F., Belmonte, C., and Ferrer-Montiel, A. (2007). A role of the transient receptor potential domain of vanilloid receptor I in channel Gating. J Neurosci 27, 11641–11650.

Goehring, A., Lee, C.H., Wang, K.H., Michel, J.C., Claxton, D.P., Baconguis, I., Althoff, T., Fischer, S., Garcia, K.C., and Gouaux, E. (2014). Screening and large-scale expression of membrane proteins in mammalian cells for structural studies. Nat Protoc 9, 2574–2585.

Gonzalez-Cobos, J.C., and Trebak, M. (2010). TRPC channels in smooth muscle cells. Front Biosci-Landmrk 15, 1023–1039.

Gregorio-Teruel, L., Valente, P., Gonzalez-Ros, J.M., Fernandez-Ballester, G., and Ferrer-Montiel, A. (2014). Mutation of I696 and W697 in the TRP box of vanilloid receptor subtype I modulates allosteric channel activation. J Gen Physiol 143, 361–375.

Grigorieff, N. (2016). Frealign: An Exploratory Tool for Single-Particle Cryo-EM. Methods Enzymol 579, 191–226.

Guo, J.T., She, J., Zeng, W.Z., Chen, Q.F., Bai, X.C., and Jiang, Y.X. (2017). Structures of the calcium-activated, non-selective cation channel TRPM4. Nature 552, 205–209.

Itsuki, K., Imai, Y., Okamura, Y., Abe, K., Inoue, R., and Mori, M.X. (2012). Voltage-sensing phosphatase reveals temporal regulation of TRPC3/C6/C7 channels by membrane phosphoinositides. Channels 6, 206–209.

Jin, P., Bulkley, D., Guo, Y.M., Zhang, W., Guo, Z.H., Huynh, W., Wu, S.P., Meltzer, S., Cheng, T., Jan, L.Y., et al. (2017). Electron cryo-microscopy structure of the mechanotransduction channel NOMPC. Nature 547, 118–122.

Kitajima, N., Numaga-Tomita, T., Watanabe, M., Kuroda, T., Nishimura, A., Miyano, K., Yasuda, S., Kuwahara, K., Sato, Y., Ide, T., et al. (2016). TRPC3 positively regulates reactive oxygen species driving maladaptive cardiac remodeling. Sci Rep-Uk 6, 1–14.

Kiyonaka, S., Kato, K., Nishida, M., Mio, K., Numaga, T., Sawaguchi, Y., Yoshida, T., Wakamori, M., Mori, E., Numata, T., et al. (2009). Selective and direct inhibition of TRPC3 channels underlies biological activities of a pyrazole compound. P Natl Acad Sci USA 106, 5400–5405.

Kumar, R., and Thompson, J.R. (2011). The Regulation of Parathyroid Hormone Secretion and Synthesis. J Am Soc Nephrol 22, 216–224.

Li, H.S., Xu, X.Z.S., and Montell, C. (1999). Activation of a TRPC3-dependent cation current through the neurotrophin BDNF. Neuron 24, 261–273.

Liu, C.H., Wang, T., Postma, M., Obukhov, A.G., Montell, C., and Hardie, R.C. (2007). In vivo identification and manipulation of the Ca2+ selectivity filter in the drosophila transient receptor potential channel. J Neurosci 27, 604–615.

Liu, X.B., Singh, B.B., and Ambudkar, I.S. (2003). TRPC1 is required for functional store-operated Ca2+ channels - Role of acidic amino acid residues in the S5-S6 region. J Biol Chem 278, 11337–11343.

Long, S.B., Tao, X., Campbell, E.B., and MacKinnon, R. (2007). Atomic structure of a voltage-dependent K+ channel in a lipid membrane-like environment. Nature 450, 376–382.

Mastronarde, D.N. (2005). Automated electron microscope tomography using robust prediction of specimen movements. J Struct Biol 152, 36–51.

Ni, C., Yan, M., Zhang, J., Cheng, R.H., Liang, J.Y., Deng, D., Wang, Z., Li, M., and Yao, Z. (2016). A novel mutation in TRPV3 gene causes atypical familial Olmsted syndrome. Sci Rep-Uk 6, 1–10.

Nilius, B., Prenen, J., Janssens, A., Owsianik, G., Wang, C.B., Zhu, M.X., and Voets, T. (2005a). The selectivity filter of the cation channel TRPM4. J Biol Chem 280, 22899–22906.

Nilius, B., Talavera, K., Owsianik, G., Prenen, J., Droogmans, G., and Voets, T. (2005b). Gating of TRP channels: a voltage connection? J Physiol-London 567, 35–44.

Oda, K., Umemura, M., Nakakaji, R., Tanaka, R., Sato, I., Nagasako, A., Oyamada, C., Baljinnyam, E., Katsumata, M., Xie, L.H., et al. (2017). Transient receptor potential cation 3 channel regulates melanoma proliferation and migration. J Physiol Sci 67, 497–505.

Ong, H.L., de Souza, L.B., and Ambudkar, I.S. (2016). Role of TRPC Channels in Store-Operated Calcium Entry. Adv Exp Med Biol 898, 87–109.

Paulsen, C.E., Armache, J.P., Gao, Y., Cheng, Y.F., and Julius, D. (2016). Structure of the TRPA1 Ion Channel Suggests Regulatory Mechanisms. Biophys J 110, 26a–26a.

Pettersen, E.F., Goddard, T.D., Huang, C.C., Couch, G.S., Greenblatt, D.M., Meng, E.C., and Ferrin, T.E. (2004). UCSF chimera - A visualization system for exploratory research and analysis. J Comput Chem 25, 1605–1612.

Prakriya, M., and Lewis, R.S. (2015). Store-Operated Calcium Channels. Physiol Rev 95, 1383–1436.

Punjani, A., Rubinstein, J.L., Fleet, D.J., and Brubaker, M.A. (2017). cryoSPARC: algorithms for rapid unsupervised cryo-EM structure determination. Nat Methods 14, 290–296.

Scheres, S.H.W. (2012). RELION: Implementation of a Bayesian approach to cryo-EM structure determination. J Struct Biol 180, 519–530.

Shen, P.S., Yang, X.Y., DeCaen, P.G., Liu, X.W., Bulkley, D., Clapham, D.E., and Cao, E.H. (2016). The Structure of the Polycystic Kidney Disease Channel PKD2 in Lipid Nanodiscs. Cell 167, 763–773.

Smyth, J.T., Hwang, S.Y., Tomita, T., DeHaven, W.I., Mercer, J.C., and Putney, J.W. (2010). Activation and regulation of store-operated calcium entry. J Cell Mol Med 14, 2337–2349.

Strubing, C., Krapivinsky, G., Krapivinsky, L., and Clapham, D.E. (2003). Formation of novel TRPC channels by complex subunit interactions in embryonic brain. J Biol Chem 278, 39014–39019.

Sudhof, T.C. (2012). Calcium Control of Neurotransmitter Release. Csh Perspect Biol 4.

Taberner, F.J., Lopez-Cordoba, A., Fernandez-Ballester, G., Korchev, Y., and Ferrer-Montiel, A. (2013). Mutations in the TRP Domain Differentially affect the Function of TRPM8. Biophys J 104, 456a–456a.

Tang, J., Lin, Y.K., Zhang, Z.M., Tikunova, S., Birnbaumer, L., and Zhu, M.X. (2001). Identification of common binding sites for calmodulin and inositol 1,4,5-trisphosphate receptors on the carboxyl termini of Trp channels. J Biol Chem 276, 21303–21310.

Vannier, B., Zhu, X., Brown, D., and Birnbaumer, L. (1998). The membrane topology of human transient receptor potential 3 as inferred from glycosylation-scanning mutagenesis and epitope immunocytochemistry. J Biol Chem 273, 8675–8679.

Vazquez, G., Wedel, B.J., Aziz, O., Trebak, M., and Putney, J.W. (2004). The mammalian TRPC cation channels. Bba-Mol Cell Res 1742, 21–36.

Winkler, P.A., Huang, Y.H., Sun, W.N., Du, J., and Lu, W. (2017). Electron cryo-microscopy structure of a human TRPM4 channel. Nature 552, 200–204.

Xia, M., Liu, D., and Yao, C. (2015). TRPC3: A New Target for Therapeutic Strategies in Chronic Pain - DAG-mediated Activation of Non-selective Cation Currents and Chronic Pain. J Neurogastroenterol 21, 445–447.

Yang, S.L., Cao, Q., Zhou, K.C., Feng, Y.J., and Wang, Y.Z. (2009). Transient receptor potential channel C3 contributes to the progression of human ovarian cancer. Oncogene 28, 1320–1328.

Yin, Y., Wu, M.Y., Zubcevic, L., Borschel, W.F., Lander, G.C., and Lee, S.Y. (2018). Structure of the cold- and menthol-sensing ion channel TRPM8. Science 359, 237–241.

Zhang, K. (2016). Gctf: Real-time CTF determination and correction. J Struct Biol 193, 1–12.

Zheng, S.Q., Palovcak, E., Armache, J.P., Verba, K.A., Cheng, Y.F., and Agard, D.A. (2017). MotionCor2: anisotropic correction of beam-induced motion for improved cryo-electron microscopy. Nat Methods 14, 331–332.

Zhu, X., Jiang, M.S., and Birnbaumer, L. (1998). Receptor-activated Ca2+ influx via human Trp3 stably expressed in human embryonic kidney (HEK)293 cells - Evidence for a non-capacitative Ca2+ entry. J Biol Chem 273, 133–142.

Zhu, X., Jiang, M.S., Peyton, M., Boulay, G., Hurst, R., Stefani, E., and Birnbaumer, L. (1996). trp, a novel mammalian gene family essential for agonist-activated capacitative Ca2+ entry. Cell 85, 661–671.

